# The use of alternative spawning habitats by the American horseshoe crab, *Limulus polyphemus*

**DOI:** 10.1101/2022.04.12.488058

**Authors:** Daniel A. Sasson, Christopher C. Chabot, Jennifer H. Mattei, Michael R. Kendrick, Jeffrey F. Brunson, Jeanette H. Huber, Jo-Marie E. Kasinak, Paul T. Puckette, Gary Sundin, Peter R. Kingsley-Smith

**Author notes:** authors contributed equally (Christopher C. Chabot, Jennifer H. Mattei). Correspondence: Daniel A. Sasson.

## Abstract

For animals that develop externally, habitats where environmental conditions are optimal for embryonic development are sometimes assumed to represent the highest recruitment potential and thus support the majority of reproductive output for a species. However, organisms may spawn in areas considered sub-optimal for embryonic development. Thus, understanding spawning habitat selection decisions and their potential impacts on recruitment and ecological interactions is necessary for predicting population status and identifying critical habitats to inform sustainable conservation decisions and effective management approaches. The American horseshoe crab, *Limulus polyphemus*, is ecologically, economically, and biomedically important. Females come ashore to spawn in the sediment where eggs develop for 2 – 4 weeks. Horseshoe crabs have been thought to primarily use sandy beach habitat for spawning in part because this habitat has been shown to be optimal for embryonic development. Horseshoe crab eggs on sandy beaches are an essential part of the diet of many organisms, including shorebirds such as the *rufa* red knot which requires the eggs to fuel their migration to arctic spawning grounds. While horseshoe crabs have been observed spawning in alternative habitats such as salt marshes and peat beds, this behavior has been assumed to be rare and non-adaptive. In this study, we compare the use of beach and alternative habitats by horseshoe crabs for spawning. To do so, we conducted adult horseshoe crab spawning surveys and horseshoe crab egg surveys in beach and *Spartina*-dominated salt marsh alternative habitats in South Carolina, Connecticut, and New Hampshire, U.S.A. While spawning horseshoe crabs were more likely to be observed on beach habitats than in alternative habitats, potentially due to logistical constraints surveying alternative habitats, we found similar densities of spawning horseshoe crabs in both habitat types. We also tended to find more eggs in alternative habitats than on beaches. Taken together, these results suggest that alternative habitats likely represent a significant source of horseshoe crab spawning activity and recruitment that had not previously been quantified. We recommend this information be incorporated into horseshoe crab population assessments, habitat protections, and more directed research at understanding variability in habitat-specific horseshoe crab spawning and its relationship to migratory shorebirds.

## Introduction

The choice of parental spawning habitat can have profound impacts on the success of offspring which may be especially true for organisms that develop externally since environmental conditions such as temperature (Gillooly and Dodson 2000; Gillooly *et al*. 2002), salinity (Przeslawski 2004; Delorme and Sewell 2014), oxygen (Cancino *et al*. 2003; Shang and Wu 2004), and pH (Findlay *et al*. 2009; Ericson *et al*. 2010) may influence development rates, viability, and recruitment to later life stages. However, spawning habitat is not always chosen based solely on embryonic developmental success and viability. For instance, spawning habitat selection may be density-dependent (Falcy 2015) or be driven by predation pressure (Korb and Linsenmair 1999), prey availability (Reglero *et al*. 2018), and anthropogenic effects (e.g., Jackson and Moser 2012). Animals may also spawn in multiple habitats as a bet-hedging strategy, a tactic adopted in heterogenous environments that seeks to maximize lifetime fitness through risk mitigation (Seger and Brockmann 1987; Lee and Doughty 2003; Simons 2009; Martin *et al*. 2019). These factors or strategies could lead animals to spawn in areas where environmental conditions appear sub-optimal for embryonic development but which provide other benefits. Determining the drivers and extent of the use of these alternative spawning habitats is necessary to predict how spawning patterns may change in the future and to understand its impact on population-level recruitment patterns.

The American horseshoe crab, *Limulus polyphemus*, is an ecologically, economically, and biomedically important species. When spawning in large numbers, they are essential bioturbators, bringing to the surface nutrients, minerals, and small invertebrates by turning over the sediment in inshore waters (Kraeuter and Fegley 1994), ultimately increasing the biodiversity and biomass of dependent species (Mattei *et al*. 2022). Their highly abundant eggs are consumed by many fish and invertebrates and are necessary fuel for many migratory shorebirds, including the rufa red knot (e.g., Keinath *et al*. 1987; Botton *et al*. 2003; Baker *et al*. 2004; Mattei *et al*. 2022). Juvenile and adult horseshoe crabs also serve as prey for multiple sea turtle species (Seney and Musick 2005, 2007). In some states, they are also harvested as bait for the eel and whelk fisheries (ASMFC 1998; Berkson and Shuster 1999; Botton 2009). Additionally, their blood is used to create Limulu*s* amoebocyte lysate (LAL), a substance that tests for the presence of bacterial endotoxins in medical devices and injectables, making LAL essential in the development and production of vaccines and critically important for pharmaceutical and medical device safety (Smith and Berkson 2005; Novitsky 2009; Krisfalusi-Gannon *et al*. 2018) For all these reasons, horseshoe crabs along the United States Atlantic coast are managed by the Atlantic States Marine Fisheries Commission (ASMFC) and individual states that harbor them.

Horseshoe crabs are traditionally thought to spawn primarily on sandy beaches (e.g., Botton *et al*. 1988; Walls *et al*. 2002; Cheng *et al*. 2021), and studies have indicated that the area around the beach high tide line provides the best conditions for embryonic development (Penn and Brockmann 1994; Vasquez *et al*. 2015a,b). Due in part to the belief that horseshoe crabs spawn primarily on sandy beaches, horseshoe crab adult spawning and egg surveys have been conducted almost entirely in beach habitats (e.g., Smith *et al*. 2002; Smith and Robinson 2015). The Adaptive Resource Management framework of the ASMFC uses in-water trawl survey data to determine the optimal level of horseshoe crab harvest in the Delaware Bay region (ASMFC 2022), with the assumption that in-water surveys provide indices of horseshoe crabs that will spawn on sandy beaches where their eggs will be available to red knots. However, horseshoe crabs have been observed to spawn in habitats other than sandy beaches in many parts of their range. For instance, in Mid-Atlantic states, horseshoe crabs sometimes spawn in muddy sediments, peat beds, and fringing salt marsh (e.g., Botton *et al*. 1988; Beekey and Mattei 2008; Castro 2019), while in New England they have been observed spawning on mudflats, and in eel grass habitats (C. Chabot, personal observation). These habitats have generally been considered unsuitable for embryonic development (e.g., Botton *et al*. 1988), but a recent study from South Carolina found that horseshoe crab clutches laid in muddy salt marsh habitat contained similar developmental stages as clutches laid on sandy beaches (Kendrick *et al*. 2021). Since egg survivorship and development can influence population growth dynamics of horseshoe crabs (Sweka *et al*. 2007; Grady and Valiela 2006), understanding the extent to which horseshoe crabs spawn in these alternative habitats will contribute to more accurate assessments of horseshoe crab populations and their conservation needs.

The goal of this study was to compare and quantify patterns of habitat use by spawning horseshoe crabs in beach habitats and alternative habitats. We chose salt marshes as the alternative habitat as this habitat is relatively abundant throughout much of the horseshoe crab geographic range (Figure 1) and previously thought to be unsuitable for embryonic development. We accomplished our goal by conducting adult horseshoe crab spawning surveys and horseshoe crab egg surveys in beach and alternative habitats in South Carolina (SC), Connecticut (CT), and New Hampshire (NH), which spans three genetically distinct populations of horseshoe crabs (King *et al*. 2015) along the United States’ Atlantic Coast. We found extensive evidence of spawning in alternative habitats for all three populations, suggesting that horseshoe crabs likely spawn regularly in non-beach habitats throughout their range.

**Figure 1.**
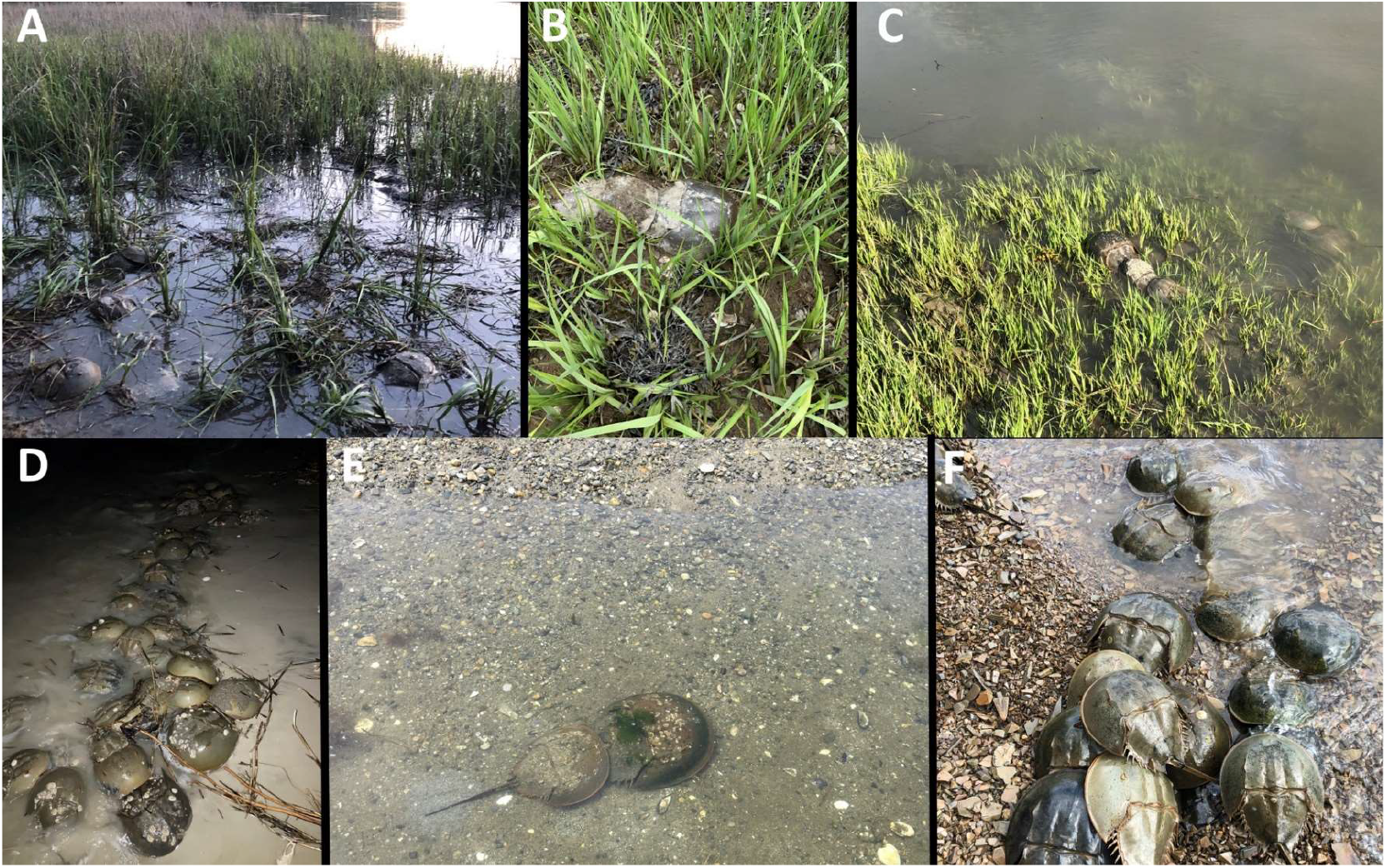
Horseshoe crabs (*Limulus polyphemus*) spawning in alternative *Spartina* dominated salt marsh in SC (A) CT (B), and NH (C), and beach habitats in SC (D), CT (E), and NH (F).

## Material and Methods

### Site Selection

In each state, we conducted adult horseshoe crab spawning surveys and horseshoe crab egg surveys at three beach habitat locations and three alternative habitat locations (Figure 2). We selected beach locations that had previously been monitored and established as horseshoe crab spawning areas, in order that high numbers of spawning adults might be expected at these locations. Conversely, many of our alternative habitat locations had not previously been verified as spawning grounds. In SC, harvesters reported collecting horseshoe crabs in salt marsh areas, but no previous spawning surveys had been conducted at our chosen locations; rather, alternative habitat locations were chosen primarily because they were near spawning beaches. In CT, alternative spawning sites were chosen based on past tagging activity tagged horseshoe crabs in marshes; however, these crabs were thought to have been caught in these areas at low tide when trying to return to the sea after spawning in nearby sandy areas. In NH, we selected alternative locations based on observations of significant horseshoe crab congregation areas from other researchers in Great Bay, but it was unclear whether spawning occurred at these locations.

**Figure 2.**
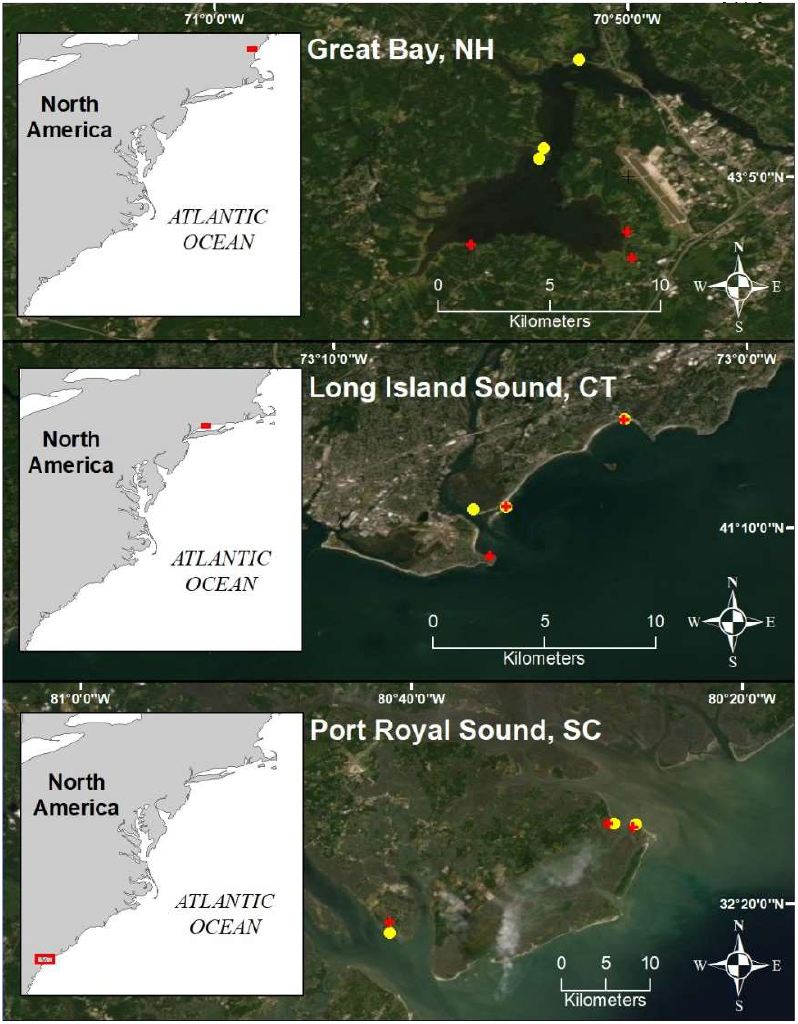
Map of survey areas. SC = South Carolina1, CT = Connecticut, NH = New Hampshire. Yellow circles indicate sandy beach habitat while red crosses indicate alternative habitats.

### Sediment temperature

We monitored the temperatures in the sediments at beach and alternative habitat locations. At each location, we buried a HOBO temperature logger (MX2203, Onset Computer Corporation, Bourne, MA) at a depth of approximately 10cm in the sediment. Temperatures were recorded every 1 (CT, NH) or 2 (SC) minutes from the start to the end of the spawning season. These start and end dates differed across states (Supplemental Table 1). We retrieved data from the HOBO loggers when conducting surveys at those locations.

### Sediment grain size

At each location, we collected single sediment cores from the beginning, middle, and end of our adult spawning and egg survey transects. In SC, we placed a randomly selected portion of each sample in a drying oven at 60°C for at least 48 hours. In CT and NH, the entire core sample was dried. After drying, we used a mortar and pestle to break apart any clumps of sediment. Each sample was then weighed to the nearest mg and poured into the top of a stack of sieves with decreasing mesh sizes (4mm, 2mm, 500μm, 250μm, 125μm, 63μm, no mesh). In SC and CT, we placed the stack of sieves on a Ro-tap sieve shaker for 15 minutes to sort the sample into its component grain size fractions. In NH, we separated the sediment in a similar manner by hand shaking for 15 minutes. We then weighed the resultant sediment remaining in each sieve to the nearest mg and calculated the percent weight of the sample for each size fraction.

### Adult horseshoe crab spawning surveys

We surveyed all locations for adult horseshoe crabs around the full and new moons in SC and CT and three to four days a week in NH since horseshoe crab spawning in Great Bay, NH is not correlated with lunar cycles (Cheng et al. 2016; see Supplemental Table 1 for locations and timing of surveys). Following protocols of Mattei (2009), the survey team at the beach walked along the transect with one observer at the high tide line and the other observer 2m into the water in SC and NH and 3m into the water in CT. For adult spawning surveys in alternative habitats in SC and NH, the survey team walked along a transect with one observer at the marsh platform edge and the other observer two meters into the marsh, away from the platform edge. It is in this area that eggs have previously been found in the salt marsh in SC (Kendrick *et al*. 2021). High turbidity and steep slopes beyond the platform edge yielded poor visibility in both states, making horseshoe crabs beyond the platform were likely uncounted. In CT, we surveyed fringing salt marshes identically to the sandy beach surveys. In all locations, if the water became murky, we used our hands and feet to feel for spawning adults. We recorded the GPS coordinates of the start and end point of each transect to determine the length and area of the transect. We recorded the sex and mating status of every adult horseshoe crab observed within the transects.

### Horseshoe crab egg surveys

We conducted egg surveys approximately one week after the spawning surveys (Supplemental Table 1). In CT and NH, we conducted one egg survey at each beach and alternative habitat location. In SC, we conducted three egg surveys at each location, one each week following the spawning surveys. At all locations, we used a mud auger (AMS One-Piece Augur) to extract sediment core samples at 40 evenly spaced points to a depth of 20cm along each spawning survey transect (transect lengths can be found in Supplemental Table 1). We then sifted through the sediment cores in the field and recorded whether eggs and/or embryos (hereafter, eggs) were found in each core sample as a binomial variable (Y/N). When present, we returned to the lab where we separated the eggs from the sediment and enumerated them. In CT and NH, we counted all eggs from each sample by hand. In SC, most samples had many thousands of eggs and so, when > approximately 1,000 eggs were present in a sample, we estimated the total number by counting the number in a randomly selected sub-sample and then used the weight of the sub-sample compared to the weight of the entire sample to estimate the total number of eggs in the sample.

### Random point horseshoe crab egg survey

In SC, between May 13, 2021 and June 14, 2021, we conducted a survey for horseshoe crab eggs at random points at beach and alternative habitat locations around Edisto Island (32.548993, - 80.297446), Pine Island (32.496871, -80.374163), and Otter Island (32.483473, -80.402966). We assigned shorelines as “marsh” or “beach” habitat and used GIS to randomly select 50 GPS points in beach habitats and 50 GPS points in marsh habitats. If we deemed a randomly selected location inaccessible in the field, we surveyed at the nearest comparable and accessible location.

Upon arrival at each location, we used hand trowels to dig to a depth of approximately 20cm, corresponding to the depth to which horseshoe crab eggs can be typically laid (Botton *et al*. 1994; Smith *et al*. 2002a), either within a few meters of the high tide line (beach locations), where horseshoe crabs are known to spawn (Penn and Brockmann 1994), or on the marsh platform where eggs have previously been found (Kendrick *et al*. 2001). We sampled at each site until we found eggs or until we had sampled for 30 person-minutes (e.g., one person sampling for 30 minutes, two people for 15 minutes, etc.), whichever occurred first. We recorded whether eggs were found at each location as a binomial variable (Y/N).

### Data analysis

To compare sediment temperature data across habitats, we calculated the daily mean, minimum and maximum temperature from each temperature logger and then calculated the mean, minimum, and maximum temperatures for all temperature loggers within a habitat type (beach or alternative) for each day, separately for each state. We then ran separate matched paired analyses for each state to compare the daily mean, minimum, and maximum temperatures across habitat types using JMP 16.1 (SAS, Cary, NC). We only included days in our analyses where we had data from at least one logger in each habitat type within a state.

To compare sediment grain size composition between habitat types, we ran generalized mixed models for each size fraction using JMP 16.1 (SAS, Cary, NC), with the size fraction percent weight as the dependent variable, habitat type as a fixed effect, and location as a random effect. We did these analyses separately for each state.

For all adult spawning and egg survey analyses, we used generalized linear mixed effects models in the lme4 and nmle packages of R v3.6.1 (Bates *et al*., 2015; R Core Team, 2019; Pinheiro *et al*. 2021). We tested model significance using chi-squared likelihood ratio tests (using the ‘anova’ function). For each analysis, fixed effects were included as well as the random effects of state, location, and location nested within state. Additionally, we compared full models with all random variables to models with reduced random variables (e.g., just state) and models with no random variables for each analysis. AIC values were used to rank models, with the lowest AIC value indicating the best fit. Below we only report statistical methods and results for the best fit model for each analysis. Code for all models can be found in the Supplemental Materials.

We analyzed our spawning survey data in two ways. First, we compared the likelihood of seeing any horseshoe crabs and then the likelihood of seeing female horseshoe crabs between beach and alternative habitats during our spawning surveys. In each model, the dependent variable was the presence of horseshoe crabs or females in the survey (Y/N), habitat type was a fixed effect and state (SC, CT, NH) was a random effect. We ran both models using the glmer function from the lme4 package (Bates *et al*. 2015; R Core Team 2019).

We then used the spawning surveys in which horseshoe crabs were seen to compare the density of spawning horseshoe crabs observed between beach and alternative habitats. Overall and female-only horseshoe crab densities were calculated for each survey by dividing the number of horseshoe crabs and females observed, respectively, by the transect area. We log-transformed overall and female-only density to normalize the data. In both models, the dependent variable was density, habitat type was a fixed effect, and survey location, state, and survey location nested within state were random effects. We ran both models using the lme function in the nmle package (Pinheiro *et al*. 2021).

We compared the likelihood of core samples from the egg surveys containing eggs between beach and alternative habitats. We ran a nominal logistic regression mixed model with the presence of eggs in each core sample (Y/N) as the dependent variable, with habitat type (beach or alternative) as a fixed effect, and survey location nested within state as a random effect. We used the glmer function from the lme4 package (Bates *et al*. 2015) for this model.

To compare egg counts between samples from the beach and alternative habitats, we used a generalized linear mixed model with a negative binomial distribution. The model contained the number of eggs in each sample as the dependent variable, habitat type as a fixed effect, and location nested within state as a random effect. For this analysis, we only included core samples in which at least one egg was found. We ran this model using the glm.nb function from the lme4 package (Bates *et al*. 2015).

We compared the likelihood of finding eggs in the beach and alternative habitats with a chi-square test, with eggs found (Y/N) as the dependent variable and habitat type (beach or alternative) as the independent variable.

## Results

### Sediment temperatures

In SC, sediments in the alternative habitat had higher mean (Supplemental Figure 1, t-ratio = -8.2, *P* < 0.001), minimum (Supplemental Figure 2, t-ratio = -6.5, *P* < 0.001), and maximum daily temperatures (Supplemental Figure 3, t-ratio = -6.1, *P* < 0.001) than sediments in the beach habitat. In CT, mean (Supplemental Figure 1, t-ratio = 8.3, *P* < 0.001), minimum (Supplemental Figure 2, t-ratio = 2.1, *P* = 0.04), and maximum (Supplemental Figure 3, t-ratio = 7.7, *P* <0.001) daily sediment temperatures were all significantly higher in beach habitats than alternative habitats. Finally, in NH, the mean (Supplemental Figure 1, t-ratio = -4.6, *P* < 0.001) and minimum (Supplemental Figure 2, t-ratio = -9.2, *P* < 0.001) daily sediment temperatures were higher in the alternative habitat than the beach habitat, but the maximum daily temperature was higher in the beach habitat than alternative habitat (Supplemental Figure 3, t-ratio = 2.1, *P* = 0.04).

### Sediment grain size analyses

In SC, grains < 63μm made up a significantly lower percentage of the sediment samples taken from sandy beach locations compared to alternative habitat locations (Supplemental Table 2). We found no significant differences in percentages for any grain size between sandy beach and alternative habitats in CT and NH (Supplemental Table 2). Location as a random effect was not significant in any of our models.

### Adult horseshoe crab spawning surveys

Overall, we saw at least one horseshoe crab in 87% (89 out of 102) of the surveys conducted at beach locations and in 55% (48 out of 87) of surveys at alternative habitat locations. We found a similar pattern when considering only females (i.e., 77% and 54%, respectively). The binomial mixed model with state as a random effect indicated that we were more likely to find horseshoe crabs at beach locations compared to alternative habitat locations (z-value = 4.6, *P* < 0.001). There was a significant effect of state as a random effect (χ^2^ = 6.4, *P* = 0.01). We found a similar effect when considering only females, i.e., they were more likely to be seen in surveys conducted at beach locations than at alternative habitat locations (z-value = 3.2, *P* < 0.01, Figure 3). We again found a significant effect of state as a random effect (χ^2^ = 3.8, *P* = 0.5).

**Figure 3.**
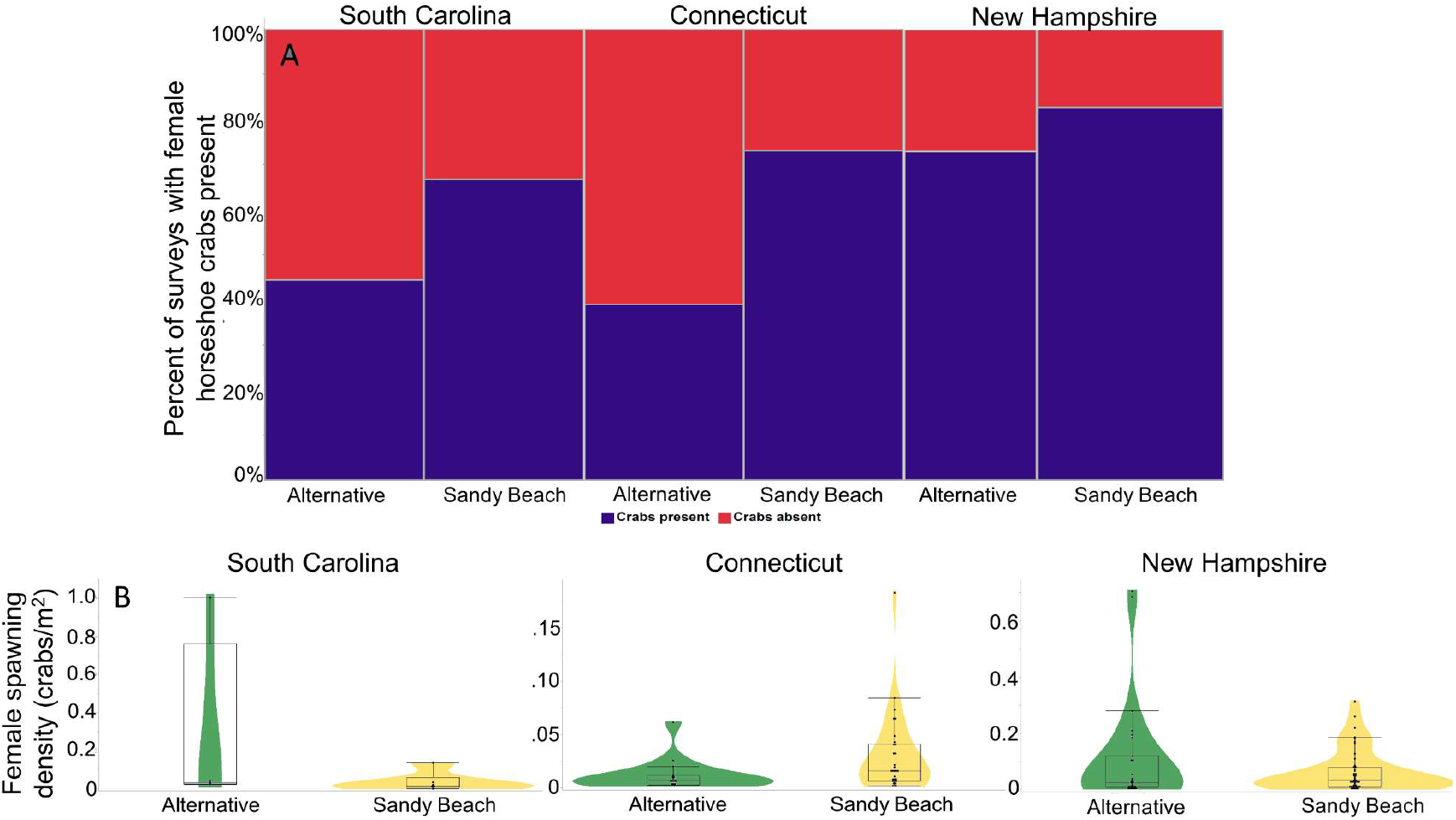
Percent of spawning surveys with females present across habitat types (A) and female spawning density in surveys with at least one female (B). Note that the scale of y-axis for female spawning density differs for each state.

We compared the density of all horseshoe crabs and females between habitat types for the surveys in which crabs were observed. We found no difference in the density of horseshoe crabs observed between habitat types (t-value = 0.30, *P* = 0.77), although there was a significant effect of location nested within state as a random effect (L. ratio = 14.0, *P* < 0.001). Female density also did not differ between habitats (t-value = 0.02, *P* = 0.99) and we again found a significant effect of survey location nested with state as a random effect (L. ratio = 15.5, *P* < 0.001).

### Horseshoe crab egg surveys

We found eggs in approximately 19% of core samples taken from beach locations and 27% of core samples taken from alternative habitat locations. Our model found no significant difference between beach and alternative habitat locations in the likelihood of finding eggs (z-ratio = 0.47, *P* = 0.64). The random effect in the model, location nested within state, was significant (χ^2^ = 109.8, *P* < 0.001). We tended to find more eggs in core samples taken from alternative locations than core samples taken from the beach locations (z-value = 1.87, *P* = 0.06). We again found that the random effect of location nested within state was significant (χ^2^ = 26.2, *P* <0.001, Figure 4).

**Figure 4.**
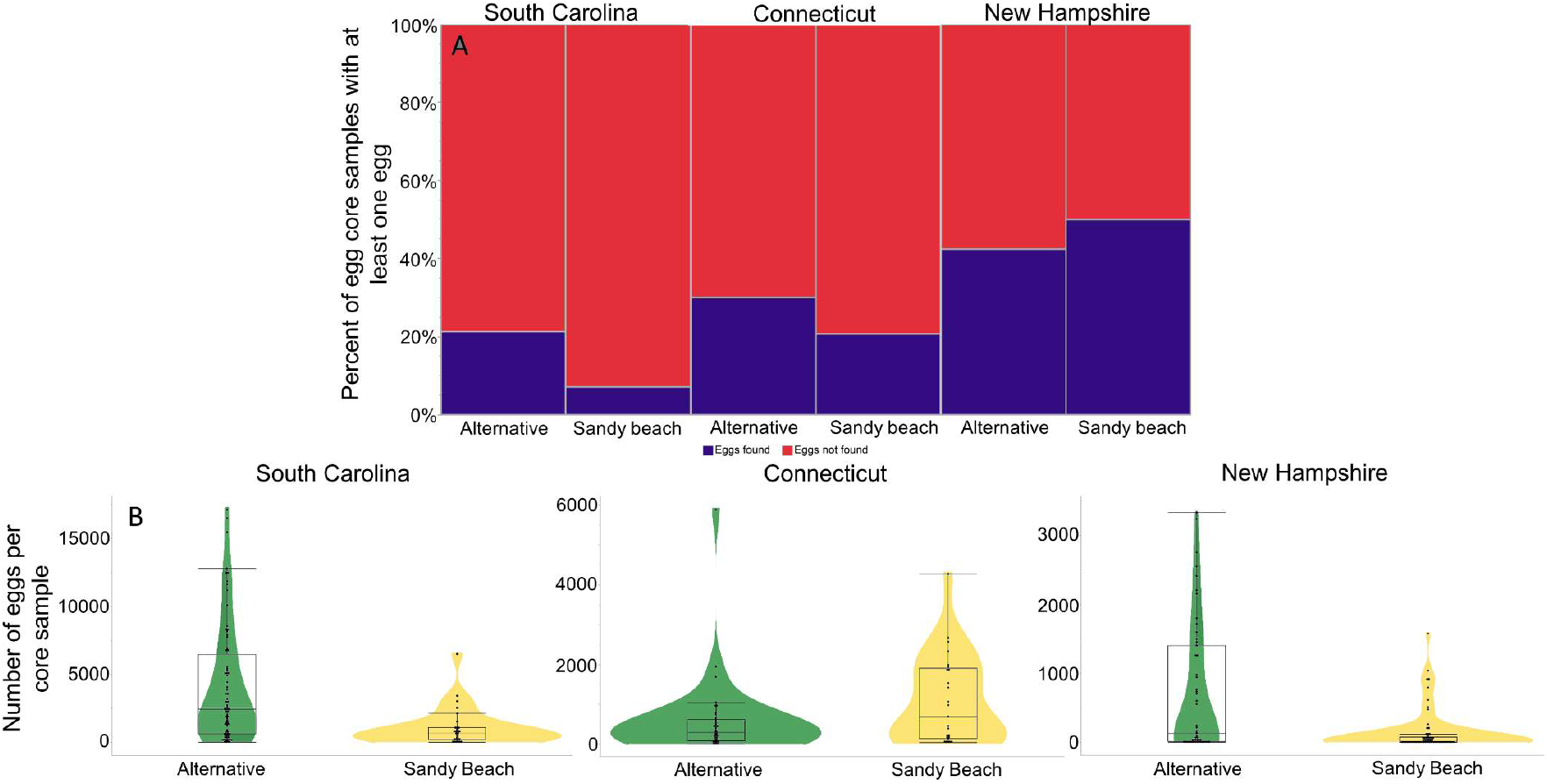
Percent of core samples from egg surveys that contained at least one egg (A) and the number of eggs in core samples that contained at least one egg (B). We tended (p = 0.06) to find more eggs in samples taken from alternative habitats than beach habitats. Note that the scale of y-axis for number of eggs differs for each state.

### Random point horseshoe crab egg survey

We found eggs at 5 out of 50 randomly selected beach locations and 7 out of 50 randomly selected alternative locations (Figure 5). There was no significant difference in the likelihood of finding eggs in either habitat (χ^2^ = 0.42, df = 1, *P* =0.51).

**Figure 5.**
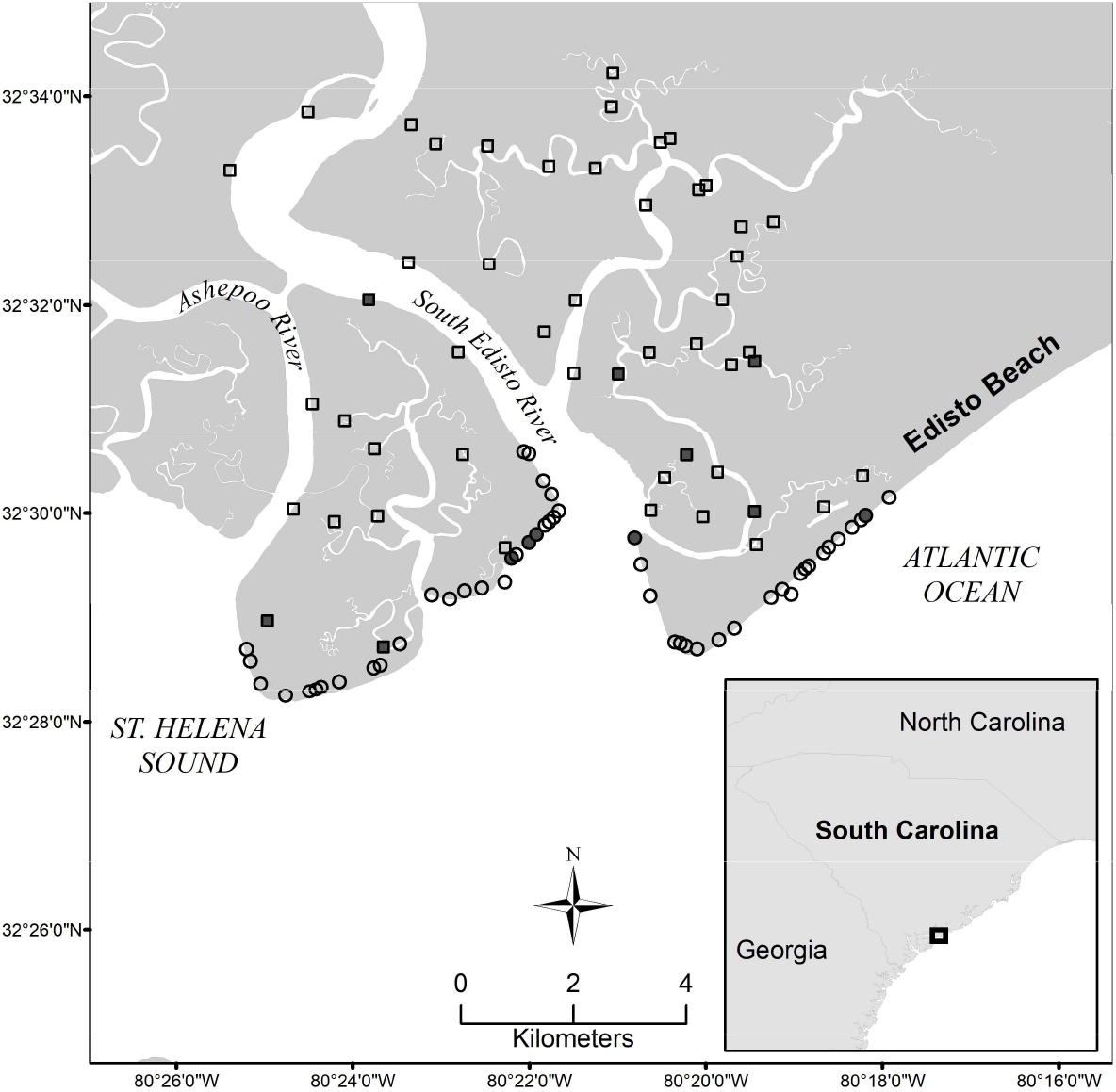
Locations of horseshoe crab egg and embryo sampling sites in the Edisto/Pine/Otter Island region within St. Helena Sound, SC. Circles represent sandy beach locations and squares alternative habitat locations. Filled in shapes indicates eggs/embryos were found at that location.

## Discussion

Our study provides evidence from three genetically distinct populations (King *et al*. 2015) that horseshoe crabs regularly spawn in habitats previously considered sub-optimal for embryonic development. Our finding supports the premise that the use of alternative spawning habitats by horseshoe crabs is geographically widespread and extensive, and thus likely an important source of recruitment for horseshoe crabs across its U.S. East Coast range. While previous studies have acknowledged some spawning by horseshoe crabs in non-sandy beach habitats (Botton *et al*. 1988; Beekey and Mattei 2008; Smith *et al*. 2017; Castro 2019), the extensive use of these habitats, based on direct observations of spawning adults, has not been previously reported. The fact that we found relatively high levels of spawning activity in alternative habitats that were somewhat haphazardly selected lends credence to our conclusion that spawning in these habitats is likely extensive. Individuals laying eggs in both beach and alternative habitats might constitute an example of bet-hedging, which can be an adaptation to temporally-or spatially-heterogenous environments that maximizes fitness over the long term through risk mitigation (Seger and Brockmann 1987; Lee and Doughty 2003; Simons 2009; Martin *et al*. 2019). In this case, individuals that nest in both beach and alternative habitats might have, on average, higher lifetime fitness than individuals that nest only in one habitat type, if there is a risk of a catastrophic loss of offspring in any given habitat due to unforeseen events (e.g., erosion, predation, nest excavations by other horseshoe crabs, etc.). However, bet-hedging can be difficult to verify (Simons 2011), and it remains unknown whether females lay eggs in both habitats, either within a season or across years, although this could be tested through mark-recapture or telemetry studies. Whether the extent of this behavior was previously undocumented or it is a relatively new phenomenon, potentially driven by a reduction in quality beach habitat, remains unclear.

Spawning surveys have been important components by which population trends are assessed (Smith *et al*. 2002b; ASMFC 2009; Beekey and Mattei 2015) but have historically been conducted almost entirely in sandy beach habitats (e.g., Smith *et al*. 2002b; Mattei *et al*. 2010; Brockmann and Johnson 2011; Smith and Robinson 2015; Cheng *et al*. 2016; Botton *et al*. 2018) and have been relied upon in some areas to estimate population or demographics trends (e.g., Smith *et al*. 2002b; Mattei *et al*. 2010; Brockmann and Johnson 2011; Beekey and Mattei 2015; Heres *et al*. 2021). Our results suggest that these surveys may be missing a large proportion of spawning adult horseshoe crabs.

Many of the alternative habitats where horseshoe crabs were found in the current study are in locations that make them logistically difficult to survey, which is a potential reason why these habitats may have been overlooked in the past. For instance, numerous potential salt marsh habitats in SC are only accessible by boat, and, during our surveys in SC, CT, and NH, the high water level on the salt marsh platform made it difficult to see spawning horseshoe crabs, increasing the likelihood of underestimating spawning horseshoe crabs in these alternative habitats. These survey challenges may explain the somewhat disconnected results from our spawning and egg surveys, although densities of adults on spawning beaches do not always directly correlate with egg abundance (Botton *et al*. 2021, but see James-Piri *et al*. 2005). For these reasons, it may be that egg surveys provide a better estimate of horseshoe crab spawning activity than spawning surveys in alternative habitats. Egg surveys may have the advantage of more directly estimating female spawning effort since not all crabs counted in spawning surveys actually spawn (e.g., Brockmann 1990). Additionally, egg surveys can be done later in the year, up to one month after the breeding season ends, and are not confined to the hours around the high tide, potentially making them logistically easier to carry out. Finally, egg surveys may be able to better account for female reproductive output in areas with low spawning densities, such as those found in CT. The appropriateness of conducting egg surveys rather than adult spawning surveys to estimate spawning activity may depend on local conditions, such as spawning density, habitat accessibility, and environmental conditions.

Previous work has shown that environmental conditions of the sediment around the high tide line of sandy beaches provide the best developmental environment for horseshoe crab embryos (Penn and Brockmann 1994; Vasquez *et al*. 2015a), and it has been suggested that some alternative habitats, such as peat bed or mud, may hinder development (Botton *et al*. 1988; Castro 2019). In SC and NH, sediment temperatures at the alternative habitat locations were generally higher than those at beach locations, while the opposite was true in CT. Higher temperatures generally increase the speed of embryonic development but can negatively impact viability when combined with low oxygen (Vasquez *et al*. 2015b). For clutches laid in SC’s muddy salt marsh habitat, embryos at all developmental stages were observed by Kendrick *et al*. (2021), although there was some indication that these eggs may have been somewhat less viable than those laid in sandy beaches. Potential lower egg viability be an indication that these sediments contain lower available oxygen than sandy beach sediments, but data is needed to test this hypothesis. Salt marsh habitat dominates much of the coastline in the southeastern U.S. (Tiner 2013) and Great Bay, NH, but are rapidly disappearing in many parts of the U.S. Atlantic Coast (Dahl 2011). Our study suggests that alternative habitats, such as salt marshes, may be a primary source of recruitment for horseshoe crabs, such that the protection of these habitats may be critical for their conservation.

One intriguing finding from our study was that we tended to find more eggs in samples taken from alternative habitats than beach habitats, with potential explanations including females laying more eggs in the alternative habitats than they do at the beach or eggs dispersing less frequently in alternative habitats. Eggs laid in sandy beaches tend to be dispersed and exhumed over time due to wave action in some areas (Nordstrom *et al*. 2006; Jackson *et al*. 2014, 2020). In SC, it may be that the generally more compact sediments, as seen by the smaller grain sizes, and lower energy waves in the alternative habitats (coastalresilience.org, 2021) tend to keep the clutches more tightly grouped. Despite grain sizes not differing significantly between beach and alternative habitats in CT and NH, and wave energy being generally similar between beach and alternative habitats in NH, we nevertheless tended to find more eggs in samples from alternative habitats than those from sandy beaches in these states as well. Egg density measurements are often used as a proxy for habitat suitability in horseshoe crabs (Smith *et al*. 2002b; Pooler *et al*. 2003; Botton *et al*. 2006, 2018, 2021). By this metric, the alternative habitats surveyed in this study are equally, if not more, suitable for horseshoe crabs than sandy beach habitats.

There may be advantages for horseshoe crabs spawning in alternative habitats. For example, one of the main causes of mortality for adult horseshoe crabs is when they are overturned by waves and cannot right themselves (Botton and Loveland 1989; Penn and Brockmann 1995). If lower energy waves or increased vegetation reduce the likelihood of overturning in some areas with alternative habitats, a reduced risk of adult mortality might outweigh any putative reduction in embryonic viability. Horseshoe crab eggs can also be an essential part of the diet for many shorebirds (e.g., Botton *et al*. 1994; Takahashi *et al*. 2019) and as well as fish and invertebrates (Botton 2009; Beekey *et al*. 2013). Predation upon these eggs may be lower in alternative habitats if, for example, shorebirds do not forage there as often or if the compactness of the sediment prevents eggs from rising to the surface. Finally, warmer sediment temperatures in alternative habitats in two states (SC and NH) could result in quicker developmental time, hatching, and metamorphosis, providing these juvenile horseshoe crabs extra time to feed and grow before their first winter.

## CONCLUSION

While beach spawning surveys are relied upon in some areas to estimate population or demographics trends (e.g., Smith *et al*. 2002b; Mattei *et al*. 2010; Brockmann and Johnson 2011; Beekey and Mattei 2015; Heres *et al*. 2021), our results suggest that these surveys may be missing much of the spawning activity by horseshoe crabs. Including alternative habitats in spawning surveys may more accurately represent population trends, although, based on our data, egg density surveys across habitats may be a better representation of reproductive effort. In addition to spawning surveys, in-water surveys are also used to assess both horseshoe crab population trends (ASMFC 2019) and to develop proxies for food availability to foraging shorebirds (ASMFC 2009). Given the extensive use of alternative habitats for spawning, the use of in-water surveys to predict egg availability to foraging shorebirds in sandy beach habitats may require further justification. Adult spawning and egg surveys on sandy beaches, however, are still valuable for understanding food availability to foraging shorebirds (e.g., Botton *et al*. 1994; Smith *et al*. 2002a; Smith *et al*. 2011). Our research highlights that while in-water trawl surveys provide valuable indices of horseshoe crab population status, constraining our understanding of horseshoe crab population dynamics to these data sources could potentially mask habitat-specific changes in ecological interactions (e.g., horseshoe crab egg availability to shorebirds). It is also unclear how climate-related factors such as sea level rise, which is increasing inundation times of salt marshes (Crotty *et al*. 2020) and leading to beach erosion in some areas (Leatherman *et al*. 2000), may lead to future changes in the relative use of sandy beach and alternative habitats by spawning horseshoe crabs.

## Acknowledgements

We would like to thank John Holloway and Marine Corp Recruit Depot for allowing us access Parris Island, SC and Keith Rossman and the community at Coffin Point, SC for providing access to the beach and salt marsh areas for surveys. We also thank the staff at the Audubon CT for access to Stratford Point, CT for surveys. We are grateful for the numerous people that helped us conduct horseshoe crab adult spawning and egg surveys, including Elizabeth Gooding, Matt King, Erica Connery at SCDNR, and undergraduate research assistants from Sacred Heart University in CT. Finally, thank you to H. Jane Brockmann who advised and commented on the grant proposal. This work was supported by a USFWS State Wildlife Grants, grant #s SC-U2-F21AP00690-00 and SC-T-F19AF00749.

