## Supplemental for "The use of alternative spawning habitats by the American horseshoe crab, *Limulus polyphemus*"

Supplemental Table 1: Locations, habitat types, transect lengths, and survey dates in the three states where research was conducted.

| Location | Habitat | Transect Length (m) | Spawning Survey Dates (2021) | Egg Survey Dates |
|---|---|---|---|---|
| *South Carolina* | | | | |
| Harbor Island | Beach | 140 - 417 | 4/25, 5/11, 5/27 | 5/3, 5/19, 6/3 |
| Harbor Island | Alternative | 50.9 - 95 | 4/25, 5/11, 5/27 | 5/3, 5/19, 6/3 |
| Coffin Point | Beach | 523 - 571 | 4/26, 5/12, 5/25 | 5/5, 5/18, 6/1 |
| Coffin Point | Alternative | 73 - 82.3 | 4/26, 5/12, 5/25 | 5/5, 5/18, 6/1 |
| Parris Island | Beach | 212 - 526.3 | 4/27, 5/10, 5/26 | 5/4, 5/17, 6/2 |
| Parris Island | Alternative | 35.4 - 99 | 4/27, 5/10, 5/26 | 5/4, 5/17, 6/2 |
| *Connecticut* | | | | |
| Milford Point Sand | Beach | 41 | 5/24, 5/26 - 5/28, 6/8, 6/10, 6/11, 6/22, 6/24, 6/26, 6/27 | 6/14 |
| Milford Point Marsh | Alternative | 142 | 5/24, 5/26 - 5/28, 6/8, 6/10, 6/11, 6/22, 6/24 - 6/27 | 6/17 |
| Milford Point Front | Beach | 154.5 | 5/24 - 5/27, 6/8, 6/10, 6/11, 6/22, 6/24, 6/26, 6/27 | 6/16 |
| Stratford Point | Alternative | 114 | 5/24, 5/26, 5/28, 6/8, 6/10, 6/11, 6/21, 6/22, 6/24 - 6/27 | 6/16 |
| Milford Harbor Sand | Beach | 31 | 5/24, 5/26, 5/27, 6/8, 6/22, 6/24, 6/26 | 6/17 |
| Milford Harbor Grass | Alternative | 33 | 5/24, 5/26, 5/27, 6/8, 6/22, 6/24, 6/26 | 6/17 |
| *New Hampshire* | | | | |
| Scammell Bridge | Beach | 60 - 95 | 5/27, 6/1 - 6/3, 6/7, 6/9.6/10. 6/15 - 6/17 | 6/17 |
| Pickering Brooke | Alternative | 100 | 6/3, 6/5, 6/7 - 6/9, 6/12, 6/14 - 6/16 | 6/17 |
| Adam's Point Western | Beach | 100 | 5/14, 5/17 - 5/19, 5/24 - 5/26, 6/1 - 6/3, 6/7 - 6/9, 6/14 - 6/17 | 6/17 |
| Great Bay Farm | Alternative | 100 | 5/12 -5/14, 5/17 - 5/19, 5/24 - 5/26, 6/1 - 6/3, 6/9, 6/14 | 6/17 |
| Adam's Boat Launch | Beach | 105 | 5/11 - 5/14, 5/17 - 5/19, 5/24 - 5/26, 5/31 - 6/3, 6/7 - 6/10, 6/14, 6/16, 6/17 | 6/17 |
| Great Bay Discovery Center - Left | Alternative | 100 | 5/17 - 5/19, 5/24 - 5/26, 5/31 - 6/3, 6/7 - 6/10, 6/14, 6/16, 6/17 | 6/17 |

Supplemental Table 2. The average percentage composition of sediment samples by grain size fractions for sandy beach and alternative habitats in each state. Bolded P-values indicate a significant difference between habitat types. -N = number of sediment samples per habitat type in each state. Location as a random effect was not significant in any models and so is not presented here.

| | *South Carolina (N = 27)* | | | *Connecticut (N = 9)* | | | *New Hampshire (N = 9)* | | |
|---|---|---|---|---|---|---|---|---|---|
| Size Fraction (μm) | Beach | Alternative | P-value | Beach | Alternative | P-value | Beach | Alternative | P-value |
| >4000 | 0.6 | 0.2 | 0.51 | 27.3 | 32.0 | 0.79 | 54.2 | 26.7 | 0.35 |
| 2000.1 – 4000 | 0.7 | 0.6 | 0.85 | 6.5 | 7.9 | 0.76 | 11.2 | 11.3 | 1.00 |
| 500.1 – 2000 | 2.6 | 7.4 | 0.08 | 35.6 | 19.6 | 0.13 | 18.5 | 29.2 | 0.48 |
| 250.1 – 500 | 9.0 | 11.4 | 0.45 | 16 | 25.1 | 0.60 | 6.7 | 14.7 | 0.15 |
| 125.1 – 250 | 75.3 | 53.4 | 0.08 | 13.8 | 14.2 | 0.96 | 5.5 | 9.6 | 0.46 |
| 63.1 – 125 | 11.0 | 14.2 | 0.62 | 0.6 | 1.0 | 0.46 | 0.9 | 8.6 | 0.10 |
| <63 | 0.9 | 12.8 | **0.04** | 0.2 | 0.2 | 0.67 | 0.0 | 0.0 | N/A |

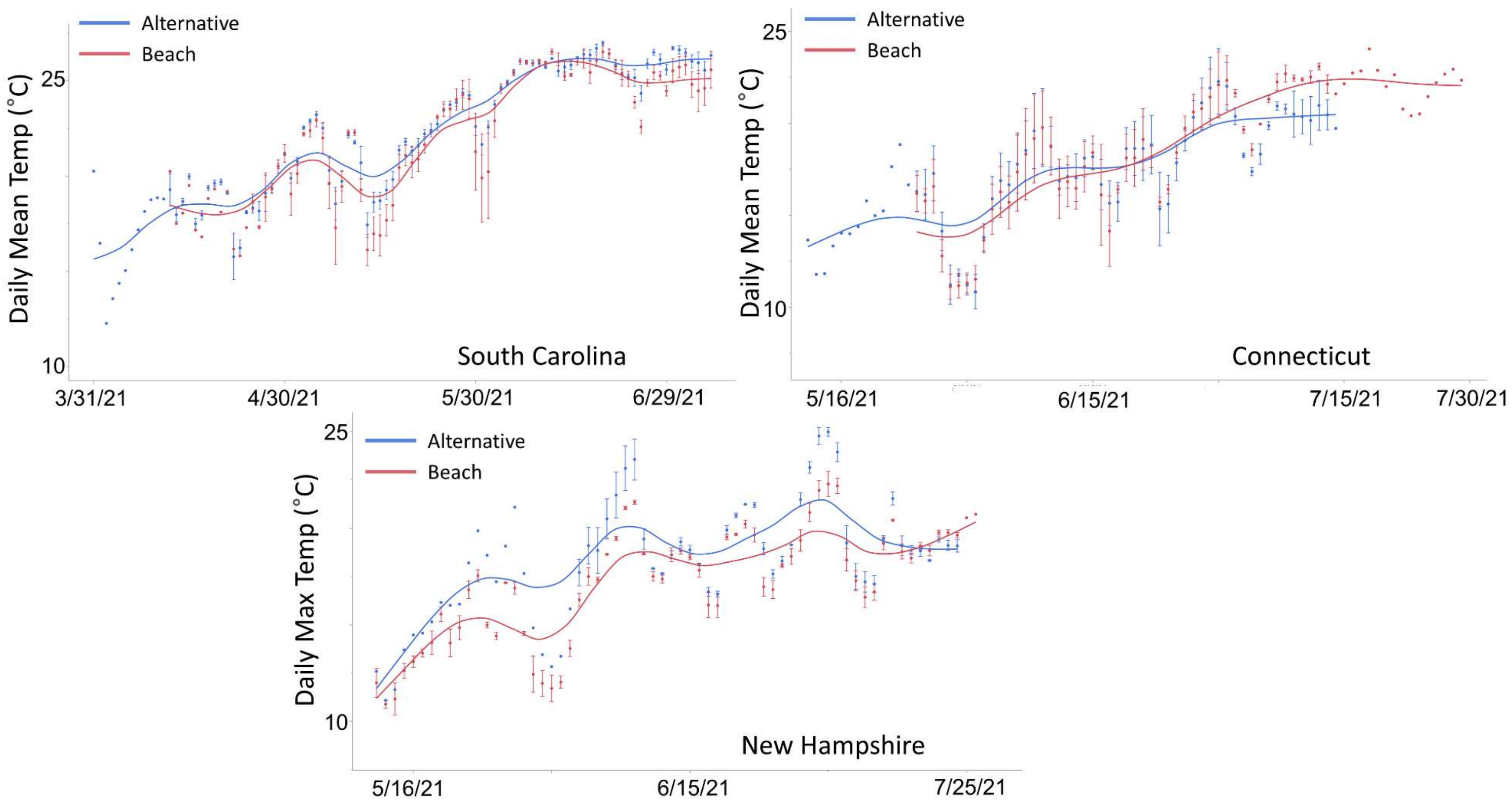


Supplemental Figure 1: Mean daily temperatures from alternative and beach sediments in South Carolina, Connecticut, and New Hampshire.

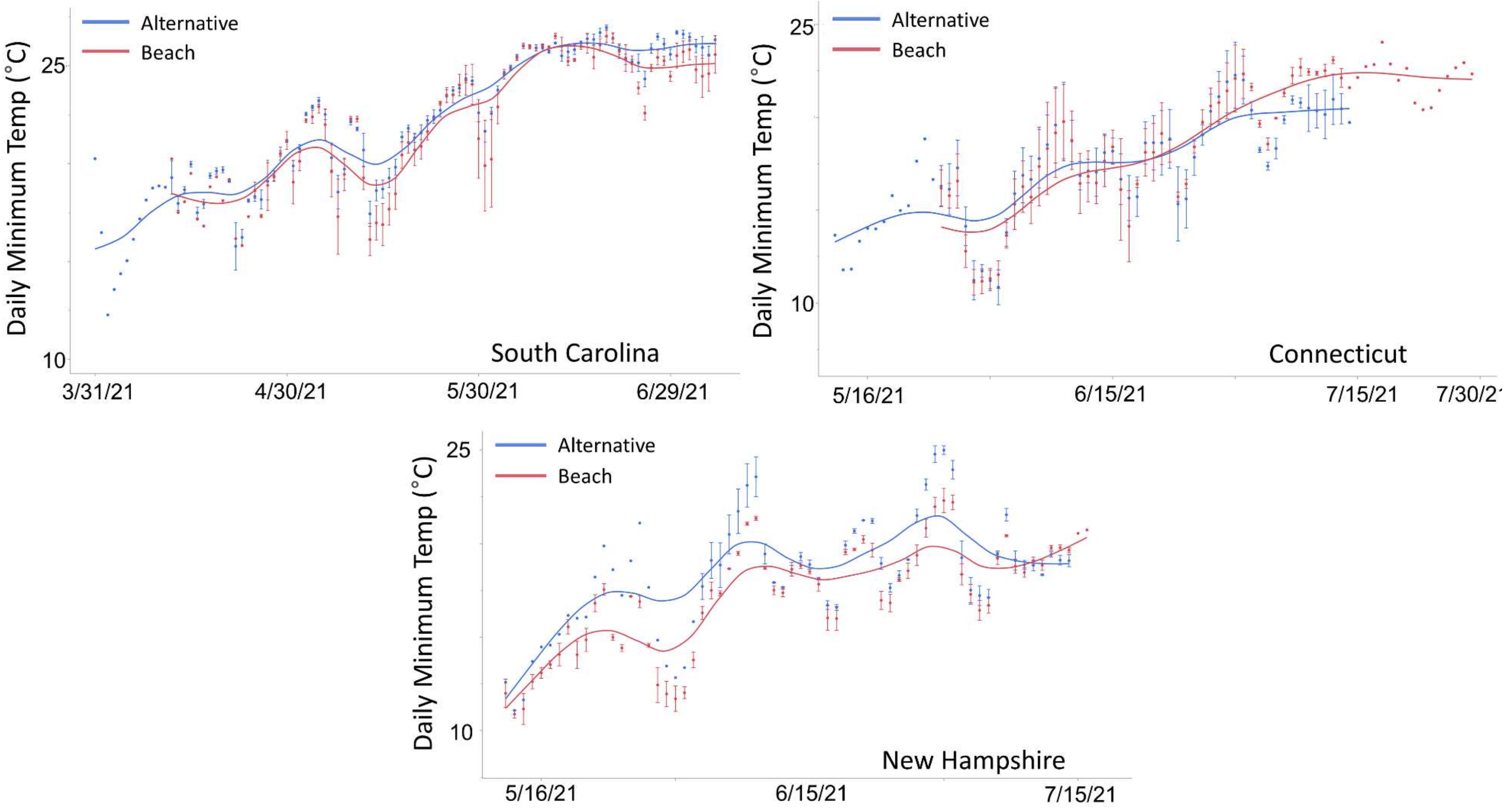


Supplemental Figure 2: Minimum daily temperatures from alternative and beach sediments in South Carolina, Connecticut, and New Hampshire.

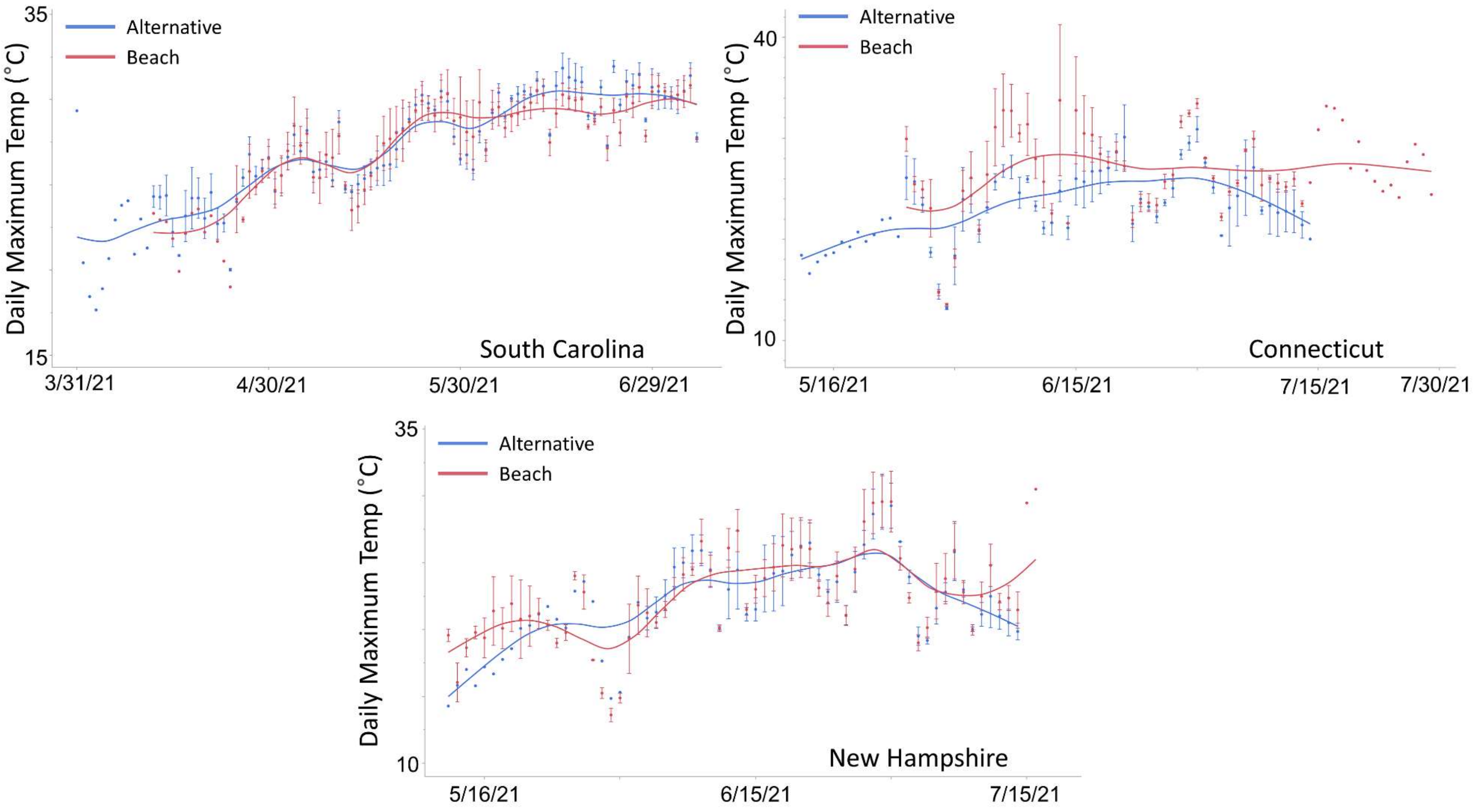


Supplemental Figure 3: Maximum daily temperatures from alternative and beach sediments in South Carolina, Connecticut, and New Hampshire.

### All and female crabs present code

```
setwd("~/horseshoe crabs/C-SWG/data/2021/For analyses")
require(ggplot2)
require(GGally)
require(reshape2)
require(lme4)
require(compiler)
require(parallel)
require(boot)
require(lattice)
require(emmeans)
require(nlme)
require(lsmeans)

overdisp_fun <- function(model) {
  rdf <- df.residual(model)
  rp <- residuals(model,type="pearson")
  Pearson.chisq <- sum(rp^2)
  prat <- Pearson.chisq/rdf
  pval <- pchisq(Pearson.chisq, df=rdf, lower.tail=FALSE)
  c(chisq=Pearson.chisq,ratio=prat,rdf=rdf,p=pval)
}

spawning_data <- read.csv("spawning_surveys_all.csv")

site <- as.factor(spawning_data$Location)
state <- as.factor(spawning_data$State)
habitat <- as.factor(spawning_data$Habitat)
total_crabs <- as.numeric(spawning_data$Total.Crabs)
females <- as.numeric(spawning_data$Female..Crabs)
total_density <- as.numeric(spawning_data$Total.Density)
female_density <- as.numeric(spawning_data$Female.Density)
density_plus <- as.numeric(spawning_data$Female.density.plus.0.0001)
femalespresent <- as.factor(spawning_data$Female.crabs.)
anycrabspresent <- as.factor(spawning_data$Any.crabs.)


#models to test for the presence of females###

model_females_full<- glmer(femalespresent ~ habitat + (1|state/site), data =  spawning_data,
family = binomial)
```

```
model_females_state <-glmer(femalespresent ~ habitat + (1|state), data = spawning_data, family = binomial)

model_females_no_random <-glm(femalespresent ~ habitat, data = spawning_data, family = binomial)

##comparing females present models###

anova(model_females_full, model_females_state, model_females_no_random)

summary(model_females_state)

anova(model_females_state, model_females_no_random)
anova(model_females_state, model_females_full)
anova(model_females_full, model_females_no_random)

females_state_pairwise <- lsmeans(model_females_state, pairwise~state, adjust = "tukey")


#models for any crabs present##

model_any_full <- glmer(anycrabspresent ~ habitat + (1|state/site), data =  spawning_data, family = binomial)

model_any_state <-glmer(anycrabspresent ~ habitat + (1|state), data = spawning_data, family = binomial)

model_any_no_random <-glm(anycrabspresent ~ habitat, data = spawning_data, family = binomial)

##comparing models##

anova(model_any_full, model_any_state, model_any_no_random)

#state model is best##

summary(model_any_state)
```

#### All crabs density code

```
setwd("~/horseshoe crabs/C-SWG/data/2021/For analyses")
require(ggplot2)
require(GGally)
require(reshape2)
require(lme4)
require(compiler)
require(parallel)
require(boot)
require(lattice)
require(emmeans)
require(nlme)

overdisp_fun <- function(model) {
  rdf <- df.residual(model)
  rp <- residuals(model,type="pearson")
  Pearson.chisq <- sum(rp^2)
  prat <- Pearson.chisq/rdf
  pval <- pchisq(Pearson.chisq, df=rdf, lower.tail=FALSE)
  c(chisq=Pearson.chisq,ratio=prat,rdf=rdf,p=pval)
}

spawning_data <- read.csv("spawning_total_present.csv")

site <- as.factor(spawning_data$Location)
state <- as.factor(spawning_data$State)
habitat <- as.factor(spawning_data$Habitat)
total_crabs <- as.numeric(spawning_data$Total.Crabs)
density_logged <-as.numeric(spawning_data$Log.Total.Crab.Density)


###model with site nested within state as a random effect###
model_normal_full2 <-lme(density_logged ~ habitat, random = (~1|state/site), data =
spawning_data)

###model with just state as random###
model_normal_state2 <- lme(density_logged ~habitat, random = ~1|state, data = spawning_data)

###model with no random effects###
model_normal_no_random <- lm(density_logged ~ habitat, data = spawning_data)
```

anova(model_normal_full2, model_normal_state2, model_normal_no_random)

##use model_normal_full2##

summary(model_normal_full2)

#### Female density code

```
setwd("~/horseshoe crabs/C-SWG/data/2021/For analyses")
require(ggplot2)
require(GGally)
require(reshape2)
require(lme4)
require(compiler)
require(parallel)
require(boot)
require(lattice)
require(emmeans)
require(nlme)

overdisp_fun <- function(model) {
  rdf <- df.residual(model)
  rp <- residuals(model,type="pearson")
  Pearson.chisq <- sum(rp^2)
  prat <- Pearson.chisq/rdf
  pval <- pchisq(Pearson.chisq, df=rdf, lower.tail=FALSE)
  c(chisq=Pearson.chisq,ratio=prat,rdf=rdf,p=pval)
}

spawning_data <- read.csv("spawning_survey_present.csv")

site <- as.factor(spawning_data$Location)
state <- as.factor(spawning_data$State)
habitat <- as.factor(spawning_data$Habitat)
total_crabs <- as.numeric(spawning_data$Total.Crabs)
females <- as.numeric(spawning_data$Female..Crabs)
total_density <- as.numeric(spawning_data$Total.Density)
female_density <- as.numeric(spawning_data$Female.Density)
density_plus <- as.numeric(spawning_data$Female.density.plus.0.0001)
femalespresent <- as.factor(spawning_data$Female.crabs.)
femalelogged <- as.numeric(spawning_data$Female.Density.logged)


###model with site nested within state as random###
model_normal_full2 <-lme(femalelogged ~ habitat, random = (~1|state/site), data =
spawning_data)

###model with just state as random###
```

```
model_normal_state2 <- lme(femalelogged ~habitat, random = ~1|state, data = spawning_data)

###mdoel with no random effects###
model_normal_no_random <- lm(femalelogged ~ habitat, data = spawning_data)

###comparing models###
anova(model_normal_full2, model_normal_state2, model_normal_no_random)

##use model_normal_full2##

summary(model_normal_full2)
```

#### Eggs found code

```
setwd("~/horseshoe crabs/C-SWG/data/2021/For analyses")
require(ggplot2)
require(GGally)
require(reshape2)
require(lme4)
require(compiler)
require(parallel)
require(boot)
require(lattice)
require(emmeans)
require(nlme)
data1 <- read.csv("All_egg_survey_all.csv")

eggsfound <-as.factor(data1$Eggs.found.)
substrate <-as.factor(data1$Beach.or.Alt)
state <-as.factor(data1$State)
site <-as.factor(data1$Location)
ID <-as.factor(data1$Sample..)


###a model with site nested within state###
model_full <- glmer(eggsfound ~ substrate + (1|state/site), data =  data1, family = binomial)

###model with just state as a random effect###
model_state <- glmer(eggsfound ~ substrate + (1|state), data = data1, family = binomial)

###Model with no random effects###
model_fixed <- glm(eggsfound ~ substrate, data = data1, family = binomial)

###comparing models###
anova(model_state, model_fixed, model_full)

anova(model_state, model_fixed, model_full)

anova(model_full, model_state)

summary(model_full)

emmeans(model_full, list(pairwise ~ substrate), adjust = "tukey")
summary(model_site)
```

### Egg numbers code

```
setwd("~/horseshoe crabs/C-SWG/data/2021/For analyses")
require(ggplot2)
require(GGally)
require(reshape2)
require(lme4)
require(compiler)
require(parallel)
require(boot)
require(lattice)
require(emmeans)
require(ggplot2)
require(sjmisc)


data1 <- read.csv("Egg_counts_all_use.csv")
substrate <- as.factor(data1$Habitat)
eggs <- as.numeric(data1$X..eggs.in.sample)
state <- as.factor(data1$State)
site <- as.factor(data1$Beach)



###Negative binomial model with site nested within state as a random effect####
eggsmodel.nested.negative <- glmer.nb(eggs ~ substrate + (1|state/site), data =  data1)
####

##State as random effect with negative binomiall##
eggsmodel.nested.negative.state <- glmer.nb(eggs ~ substrate + (1|state), data =  data1)

#negative binomial model with no random effect##
eggsmodel.no.random <-glm.nb(eggs ~ substrate, data = data1)

##comparing two negative binomial models##
anova(eggsmodel.nested.negative, eggsmodel.nested.negative.state, eggsmodel.no.random)

anova(eggsmodel.nested.negative.state, eggsmodel.nested.negative)


##full model is best##

summary(eggsmodel.nested.negative)
```

```
emmeans(eggsmodel.nested.negative, list(pairwise ~ substrate), adjust = "tukey")
```

### Random point egg survey code

```
setwd("~/horseshoe crabs/Internal SWG/data/2021")

require(ggplot2)
require(GGally)
require(reshape2)
require(lme4)
require(compiler)
require(parallel)
require(boot)
require(lattice)
require(emmeans)
require(nlme)

swg <- read.csv("SWG_data.csv")

eggs <- as.factor(swg$Eggs..Y.N.)
habitat <- as.factor(swg$Habitat)

## testing the likelihood of finding eggs in marsh and beach habitat
chisq <- chisq.test(eggs, habitat, correct = FALSE)

chisq
```
